# Spatial vascular/BTB remodeling and malignant-state plasticity in glioblastoma

**DOI:** 10.64898/2026.08.18.745357

**Authors:** Li Zheng, Lu Gan

## Abstract

**Background:** Glioblastoma (GBM) contains spatially heterogeneous malignant and vascular states, but blood-tumor barrier (BTB) remodeling is often described as a binary functional phenotype. We asked whether anatomically distinct GBM compartments contain separable vascular programs that coexist with malignant-state plasticity.

**Methods:** We performed donor-aware cross-sectional analyses of 38 histopathology-annotated spatial transcriptomic sections from 6 donors and a separately analyzed endothelial single-nucleus layer from the same GBM-Space atlas. Complementary external datasets tested patient-paired regional remodeling, anatomical replication, cross-technology source localization, and malignant-state architecture.

**Results:** *THSD1*-*FLT4* Recognition increased from leading edge to infiltrative tumor (median adjusted effect +0.02875; 4/4 donors positive). Priming increased across this boundary (+0.14814; 3/4) but decreased from infiltrative to cellular tumor (-0.16409; 0/4), whereas Gate remodeling increased from infiltrative to cellular tumor (+0.21296; 4/4). Remodeled endothelium showed higher *PLVAP* detection (+0.26409; 12/12 donors) and *PLVAP* pseudobulk expression (+1.61784 log1pCPM; 11/12), with lower *MFSD2A* pseudobulk expression (-0.71448; 10/12 negative). External cohorts supported regional vascular/BTB remodeling, while GSE131928 supported broad malignant-state architecture and an exploratory within-tumor pseudotemporal continuum.

**Conclusions:** GBM contains spatially partitioned vascular/BTB-associated programs alongside malignant-state plasticity. Recognition-Priming-Gate is a cross-sectional discovery framework, not a validated temporal cascade, and the data do not establish BTB permeability, causal tumor-vascular signaling, or therapeutic-delivery benefit.

**Key Points:**

- BTB-associated remodeling is spatially partitioned across GBM compartments.
- A *PLVAP*-high/*MFSD2A*-low endothelial state marks remodeled vasculature.
- Malignant-state plasticity forms a parallel, non-causal transcriptional axis.

**Importance of the Study:** The blood-tumor barrier in glioblastoma is often discussed as either intact or disrupted, yet therapeutic access is likely shaped by regionally distinct vascular states. This study resolves donor-aware spatial differences across leading-edge, infiltrative, and cellular-tumor compartments and identifies a *PLVAP*-high/*MFSD2A*-low endothelial remodeling state associated with the cellular-tumor region. Independent datasets support the broader vascular-remodeling phenotype, while a separate adult cohort supports malignant-state plasticity as a parallel axis. The Recognition-Priming-Gate framework is intentionally cross-sectional and does not claim a temporal cascade or measured barrier opening. By defining where barrier-associated transcriptional states are spatially organized, the study provides a testable map for future functional experiments aimed at transient, spatially targeted modulation of therapeutic delivery.

## Introduction

Glioblastoma (GBM) is a lethal primary brain malignancy characterized by marked cellular, microenvironmental, and vascular heterogeneity. The blood-tumor barrier (BTB) is likewise heterogeneous: vascular permeability and endothelial programs differ across tumor regions, and intact barrier features can persist in infiltrative tissue even when the angiogenic core is disrupted.^1,2^ A binary “open-versus-closed” description therefore incompletely captures the molecular organization of the GBM vasculature.

GBM malignant cells also occupy recurrent OPC-like, NPC-like, AC-like, and MES-like transcriptional states and can display substantial within-tumor plasticity.^3^ These state frameworks provide a biological context for spatial GBM studies, but they do not by themselves establish how vascular or BTB-associated programs are organized across anatomical compartments. Conversely, endothelial remodeling signatures should be interpreted in the malignant and stromal context in which they occur.

Spatial transcriptomics creates an additional inferential challenge. Sections, spots, cells, and nuclei provide dense measurements but are nested within donors and tumors. Treating nested observations as independent biological replicates can overstate certainty, particularly when cross-sectional regional contrasts are interpreted as temporal trajectories or mechanisms. Donor- or tumor-aware analyses are therefore needed to distinguish anatomical organization from temporal progression and transcript localization from physiological barrier function.

Here, we asked whether histopathology-annotated leading-edge (LE), infiltrative-tumor (IT), and cellular-tumor (CT) compartments resolve separable vascular/BTB-associated programs. We evaluated a *THSD1*-*FLT4*-associated Recognition score, a composition-residualized Priming state, and a Gate-remodeling state as distinct regional behaviors, explicitly treating Recognition-Priming-Gate as a cross-sectional discovery framework rather than a validated temporal sequence. We further examined whether endothelial transcriptional remodeling and malignant-cell state plasticity form two structured but non-causal axes and tested defined components of the model across complementary external datasets.

## Materials and Methods

### Study design and data resources

We performed a secondary, cross-sectional analysis of de-identified, previously generated transcriptomic datasets. The primary resource was the public GBM-Space multimodal atlas, which contains pathology-annotated spatial transcriptomic data and a separately analyzed single-nucleus layer.^4^ The present spatial analysis used 38 sections from 6 donors with the required histopathology, cell-state abundance, spatial-niche abundance, and gene-expression features. The single-nucleus object contained 1,025,329 nuclei and 36,601 features from 12 tumors; this layer belongs to the same GBM-Space atlas and was not treated as an independent external cohort. No new participant recruitment, tissue collection, or intervention was performed.

### Spatial compartments and eligibility

Histopathology annotations were collapsed into leading-edge (LE), infiltrative-tumor (IT), cellular-tumor (CT), vascular, and necrotic compartments. LE, IT, and CT were treated as cross-sectional anatomical labels. High-confidence LE/IT/CT assignment required exactly one regional score to be at least 0.50. The primary complete-triad MIN50 cohort required at least 50 high-confidence locations in each LE, IT, and CT region and retained 5 sections from 4 donors; the IT-CT MIN50 cohort retained 17 sections from the same 4 donors. MIN25 selected the same sections as MIN50, and a prespecified MIN100 sensitivity analysis retained 4 complete sections, one from each donor. Sections and spots quantified sampling depth and were not treated as independent patients (Supplementary Fig. S1; Supplementary Table S1).

### Expression and module analysis

Gene-expression counts were normalized by full Gene Expression library size and log transformed. Within each section, genes were standardized across the analyzed compartments, and prespecified BTB/vascular modules were calculated as arithmetic means of component-gene Z scores. Regional means were estimated within section, section contrasts were calculated as IT minus LE or CT minus IT, and multiple sections from the same donor were collapsed by their median before across-donor summaries. Threshold and model robustness are summarized in Supplementary Fig. S2 and Supplementary Table S2.

### Recognition, Priming, and Gate

The Recognition analysis focused on *THSD1*, *FLT4*, and *MPZL3*. The *THSD1*-*FLT4* pair score was the minimum of the two within-section Z scores, and the *THSD1*-*FLT4*-*MPZL3* co-state score was the minimum of all three Z scores; these minimum-based scores require relative co-elevation at a spatial location but do not establish same-cell expression or biochemical interaction. Ordinary least-squares models were fit within section with regional indicator, standardized vascular abundance, and malignant fraction as covariates, then collapsed to donor medians. For the continuous-state analysis, the THSD1-FLT4-MPZL3 co-state score served as the Recognition/LOCK term. The LOCK, transcytosis-core, and junctional-barrier scores were each residualized within section against an intercept, standardized vascular abundance, and standardized malignant fraction; the residuals were re-standardized within section to obtain Z_LOCK_RESID, Z_TRANS_RESID, and Z_JUNCTION_RESID. The continuous Priming score was (Z_LOCK_RESID + Z_JUNCTION_RESID - Z_TRANS_RESID)/3, and the Gate-conversion score was (Z_TRANS_RESID - Z_JUNCTION_RESID)/2. Prespecified quantile sensitivity analyses used thresholds from 0.50 to 0.70.

### Supplementary donor and spatial-topology analyses

Primary effects were summarized at the biological-unit level in a prespecified supplementary robustness analysis, with exact sign tests and biological-unit bootstrap intervals treated descriptively (Supplementary Fig. S3; Supplementary Table S3). In the same five-section Visium discovery cohort, *THSD1*-positive malignant-enriched spots were prespecified as source niches and *FLT4*-positive vascular-enriched spots as target niches. Same-spot co-occurrence, neighborhood enrichment, and nearest-target distance were tested against within-section LE/IT/CT-stratified permutations; section estimates were aggregated within donor. The primary q50 analysis used 5,000 permutations and q60/q70 sensitivity analyses used 2,000 permutations. This analysis tests spatial association, not direct cell-cell contact or signaling (Supplementary Fig. S4; Supplementary Table S4).

### Endothelial-state and transcript-source analyses

Endothelial nuclei were selected from the GBM-Space single-nucleus layer and grouped as canonical endothelium (capillary, arteriole, venule; *n* = 1,363) or remodeled endothelium (CNA-associated and other endothelial populations; *n* = 878), for 2,241 endothelial nuclei from 12 donors. *PLVAP* detection and donor-level pseudobulk expression of *PLVAP* and *MFSD2A* were compared between states. Pseudobulk counts were normalized to log(1+CPM); biological donors, rather than nuclei, were resampled in the biological-unit bootstrap analysis. Separately, transcript-source localization compared *MFSD2A*, *PLVAP*, *CAV1*, and *CAVIN1* across 9 prespecified vascular, mural, and other source groups, preserving donor as the inferential unit. Fine vascular-composition models were retained as sensitivity analyses; early IT-versus-LE detailed models were interpreted with collinearity caution.

### External validation

GSE162631 was used for patient-paired regional endothelial remodeling and was analyzed at the patient level.^5^ The corrected Ivy Glioblastoma Atlas Project/GSE107559 ASTR122 cohort was used for tumor-level anatomical replication across LE, IT, and CT, aggregating repeated regional samples within tumor.^6^ Strict source-annotated Xenium S2/S6 data were used only for cross-technology transcript-source localization.^7^ GSE131928 was used for independent adult malignant-state architecture and within-tumor pseudotemporal organization using author-defined OPC-, NPC-, AC-, and MES-associated scores.^3^ These datasets were assigned non-overlapping evidentiary roles and were not pooled into a single statistical cohort (Supplementary Fig. S5; Supplementary Tables S5-S6).

### Statistical principles and software

Donor, patient, or tumor was the biological inferential unit appropriate to each dataset. Directional consistency, effect sizes, and biological-unit summaries were emphasized, particularly where the number of independent spatial donors was small; 4-of-4 direction consistency corresponds to a two-sided exact sign-test *P* = 0.125 and was not described as conventional statistical significance. Analyses used 64-bit Microsoft Windows, Python 3.11.15, JupyterLab 4.6.2, Scanpy 1.11.5, AnnData 0.12.19, h5py 3.16, NumPy 2.4.6, pandas 2.3.3, SciPy 1.17.1, statsmodels 0.14.6, Matplotlib 3.11.1, seaborn 0.13.2, and scikit-learn 1.9.0. Additional reproducibility details are provided in the Supplementary Methods.

### Generative AI assistance

The authors used OpenAI’s ChatGPT (version GPT-5.6, accessed August 2026) solely to assist with language editing and improve the readability of the manuscript. All scientific judgments, interpretations, and conclusions were independently made by the authors. The authors take full responsibility for all scientific content of the manuscript.

## Results

### A donor-anchored anatomical framework defines the spatial cohort

The discovery cohort comprised 38 histopathology-annotated sections from 6 donors across LE, IT, CT, vascular, and necrotic compartments. Seventeen sections from 4 donors were eligible for the IT-CT MIN50 comparison; 5 sections from the same 4 donors formed complete LE-IT-CT MIN50 triads, and 4 complete sections from the same donors remained at MIN100. The cohort and module tables contained 190 section-by-region and 135 section-by-region-by-module records, respectively, but these records represented nested sampling rather than independent biological replicates (Fig. 1; Supplementary Fig. S1; Supplementary Table S1).

**Figure 1.**
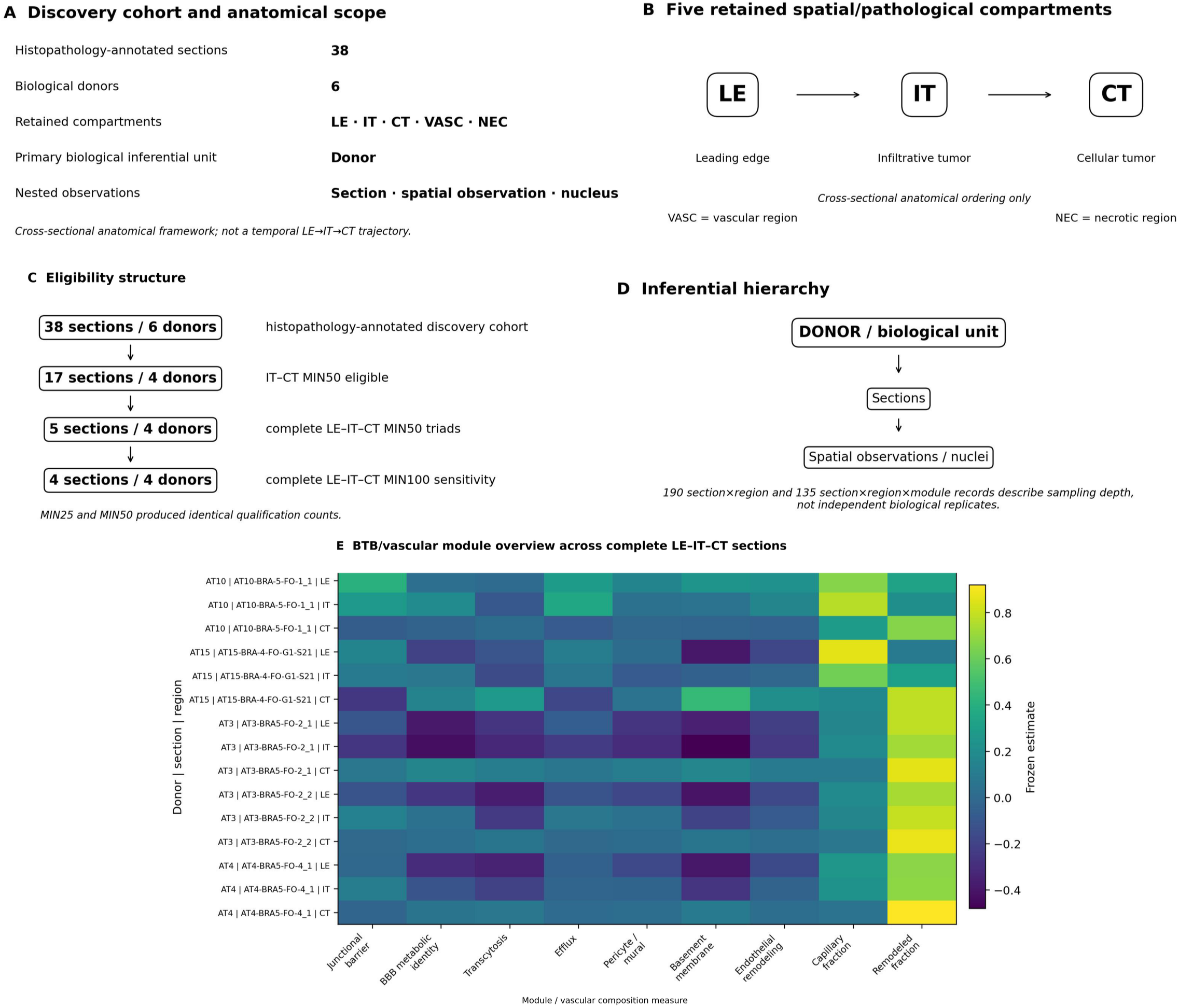
Cohort structure and the cross-sectional LE-IT-CT anatomical framework. (A) Overview of the discovery cohort comprising 38 histopathology-annotated sections from 6 donors. (B) Schematic of the five retained compartments, with LE, IT, and CT shown as a cross-sectional anatomical ordering and VASC and NEC retained as additional pathological compartments. (C) Eligibility summary showing 17 IT-CT MIN50-eligible sections from 4 donors, 5 complete LE-IT-CT MIN50 triads from 4 donors, and 4 complete triads retained under MIN100; MIN25 and MIN50 produced identical qualification counts. (D) Donor-anchored sampling structure distinguishing biological donors from nested sections and spatial observations. (E) Source-traceable spatial overview of BTB/vascular modules across complete three-zone sections. The framework is anatomical and cross-sectional and does not imply temporal LE-to-IT-to-CT progression. **Alt text:** Cohort and spatial maps show 38 sections from six donors, LE/IT/CT/VASC/NEC compartments, eligibility thresholds, and donor-nested sampling.

### *THSD1-FLT4* Recognition and Priming are regionally distinct from Gate remodeling

The adjusted *THSD1*-*FLT4* Recognition score increased from LE to IT (median donor effect, +0.02875; 4/4 donors positive) but was mixed from IT to CT (-0.01621; 2/4 positive). The *THSD1*-*FLT4*-*MPZL3* score showed a concordant LE-to-IT increase (+0.01499; 4/4 positive), whereas the incremental *MPZL3* contrast was heterogeneous (+0.02334; 2/4 positive). Priming increased from LE to IT (+0.14814; 3/4 positive) and decreased from IT to CT (-0.16409; 0/4 positive). Gate showed the complementary pattern, decreasing from LE to IT (-0.06408; 1/4 positive) and increasing from IT to CT (+0.21296; 4/4 positive) (Fig. 2). In the donor-level decomposition, *THSD1* showed the expected LE-to-IT direction in 4/4 donors and *FLT4* in 3/4, while the composite Recognition score remained positive in 4/4 (Supplementary Fig. S3; Supplementary Table S3).

**Figure 2.**
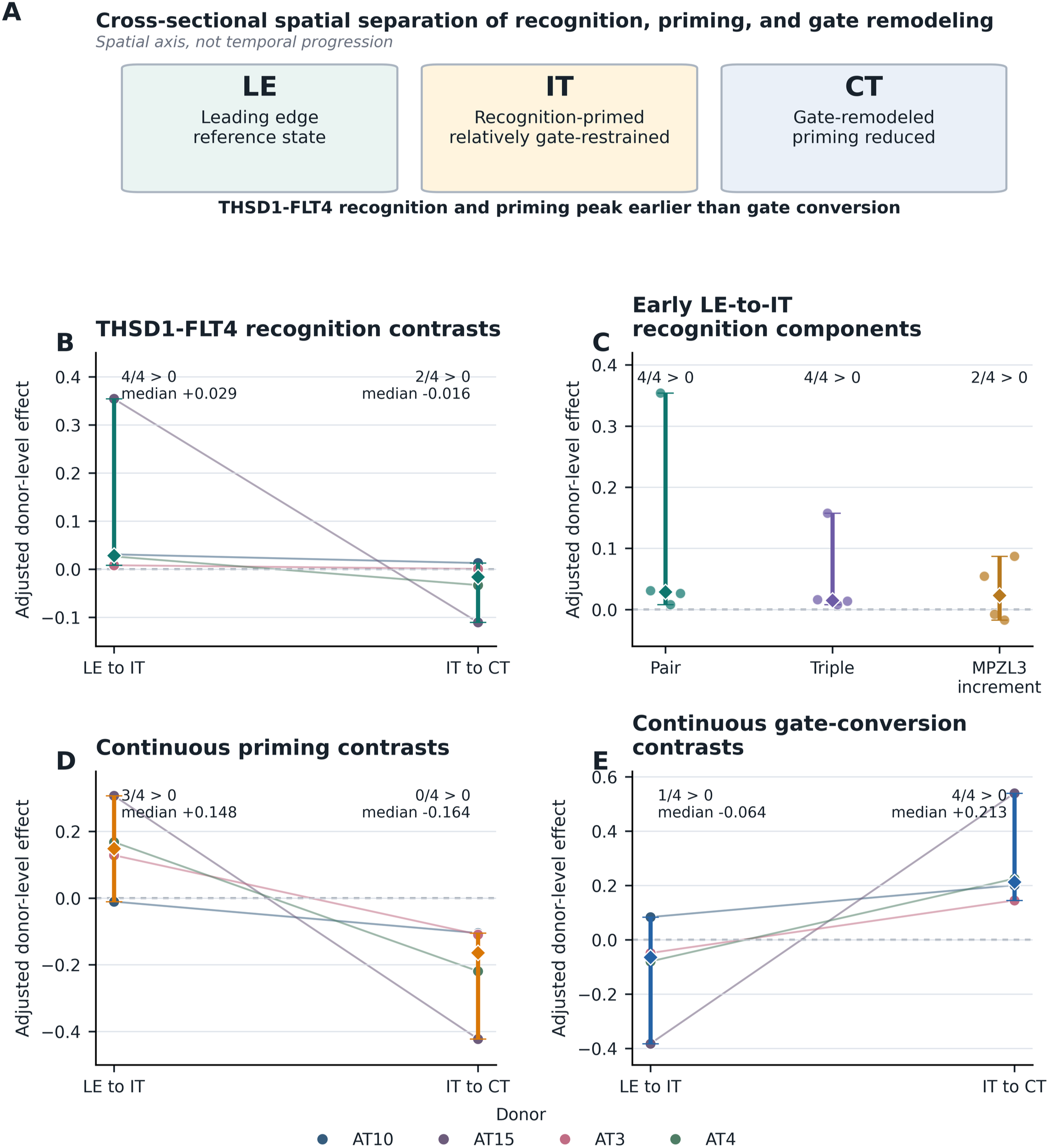
*THSD1*-*FLT4* Recognition, Priming, and Gate remodeling across the cross-sectional LE-IT-CT anatomical axis. (A) *THSD1*-*FLT4* Recognition: LE-to-IT median adjusted effect +0.02875 with 4/4 positive donors; IT-to-CT -0.01621 with 2/4 positive donors. (B) *THSD1*-*FLT4*-*MPZL3* score: LE-to-IT +0.01499 with 4/4 positive donors; *MPZL3* incremental contrast +0.02334 with 2/4 positive donors. (C) Priming: LE-to-IT +0.14814 with 3/4 positive donors and IT-to-CT -0.16409 with 0/4 positive donors. (D) Gate conversion: LE-to-IT -0.06408 with 1/4 positive donors and IT-to-CT +0.21296 with 4/4 positive donors. (E) Integrated donor-direction summary. The figure represents cross-sectional regional organization and does not establish a temporal sequence, biochemical AND gate, permeability change, or causal coupling. **Alt text:** Donor-level plots show Recognition and Priming concentrated at LE-to-IT and Gate remodeling concentrated at IT-to-CT, with direction counts.

### *THSD1*-*FLT4* spatial topology supports association beyond regional structure

A prespecified same-cohort topology analysis further supported spatial association between the Recognition components. *THSD1*-positive malignant-enriched niches showed positive neighborhood enrichment with *FLT4*-positive vascular-enriched niches in 4/4 donors after LE/IT/CT-stratified permutation. Same-spot co-occurrence and nearest-target closeness were positive in 3/4 donors, and direction patterns were retained across q50, q60, and q70 sensitivity thresholds. These results support spatial proximity beyond regional anatomical structure but not direct tumor-endothelial contact or biochemical *THSD1*-*FLT4* signaling (Supplementary Fig. S4; Supplementary Table S4).

### A *PLVAP*-high/*MFSD2A*-low endothelial axis characterizes remodeled endothelium

The single-nucleus endothelial layer contained 1,363 canonical and 878 remodeled endothelial nuclei from 12 donors. *PLVAP* detection was higher in remodeled than canonical endothelium by +0.26409, with 12/12 donors positive. *PLVAP* pseudobulk expression was also higher (+1.61784 log1pCPM; 11/12 positive), whereas *MFSD2A* pseudobulk expression was lower (-0.71448 log1pCPM; 10/12 negative) (Fig. 3; Supplementary Fig. S3; Supplementary Table S3). These measurements define a donor-reproducible transcriptional state; they are not a permeability assay.

**Figure 3.**
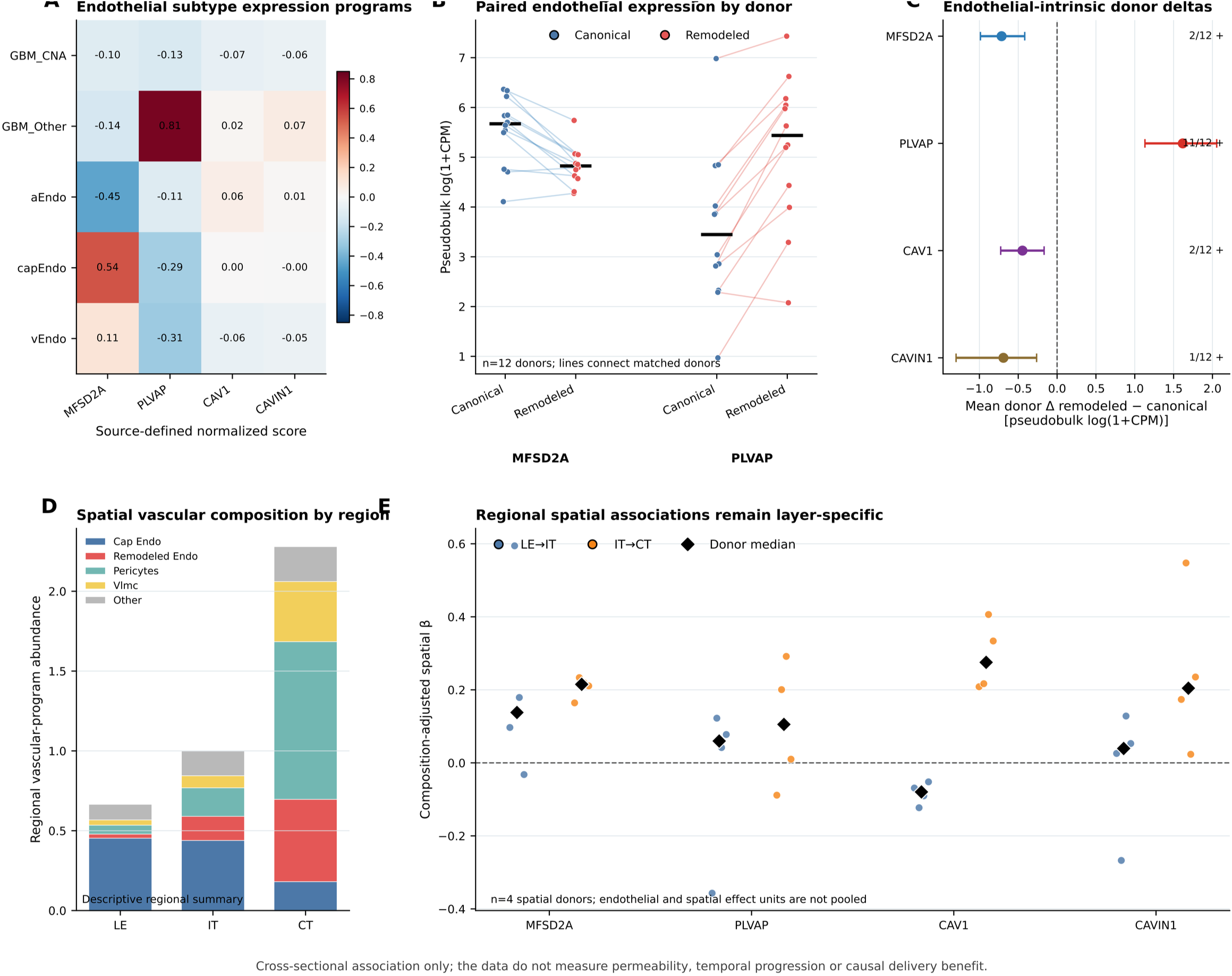
Donor-aware endothelial and spatial composition evidence for the remodeled Gate-associated vascular state. (A) Source-defined normalized scores for *MFSD2A*, *PLVAP*, *CAV1*, and *CAVIN1* across endothelial subtypes. (B) Paired canonical-versus-remodeled endothelial pseudobulk for *MFSD2A* and *PLVAP* in 12 donors. (C) Mean remodeled-minus-canonical donor deltas for all four genes, with displayed intervals and positive-donor counts. (D) Descriptive vascular-program composition across LE, IT, and CT. (E) Donor-level composition-adjusted LE-to-IT and IT-to-CT spatial associations for all four genes. The figure supports a *PLVAP*-high/*MFSD2A*-low endothelial transcriptional remodeling axis but does not establish BTB permeability, barrier opening, protein function, or causal remodeling. **Alt text:** Endothelial analyses compare canonical and remodeled states and show higher *PLVAP* and lower *MFSD2A* in remodeled endothelium across donors.

### Transcript-source localization separates endothelial anchors from distributed vascular signals

Canonical endothelium was the top *MFSD2A* source in 9/9 complete donors (median endothelial advantage, +0.9237 log1pCPM), whereas remodeled endothelium was the top *PLVAP* source in 8/9 donors (+2.1375 log1pCPM). *CAV1* had a distributed source architecture: pericytes were the most frequent top source in only 3/9 donors, the median endothelial advantage was -0.052 log1pCPM, and its composition-adjusted IT-to-CT spatial effect was +0.27529 with 4/4 positive spatial donors. *CAVIN1* was predominantly canonical-endothelial in 7/9 donors (+0.5157 log1pCPM) but was shared with pericytes (Fig. 4).

**Figure 4.**
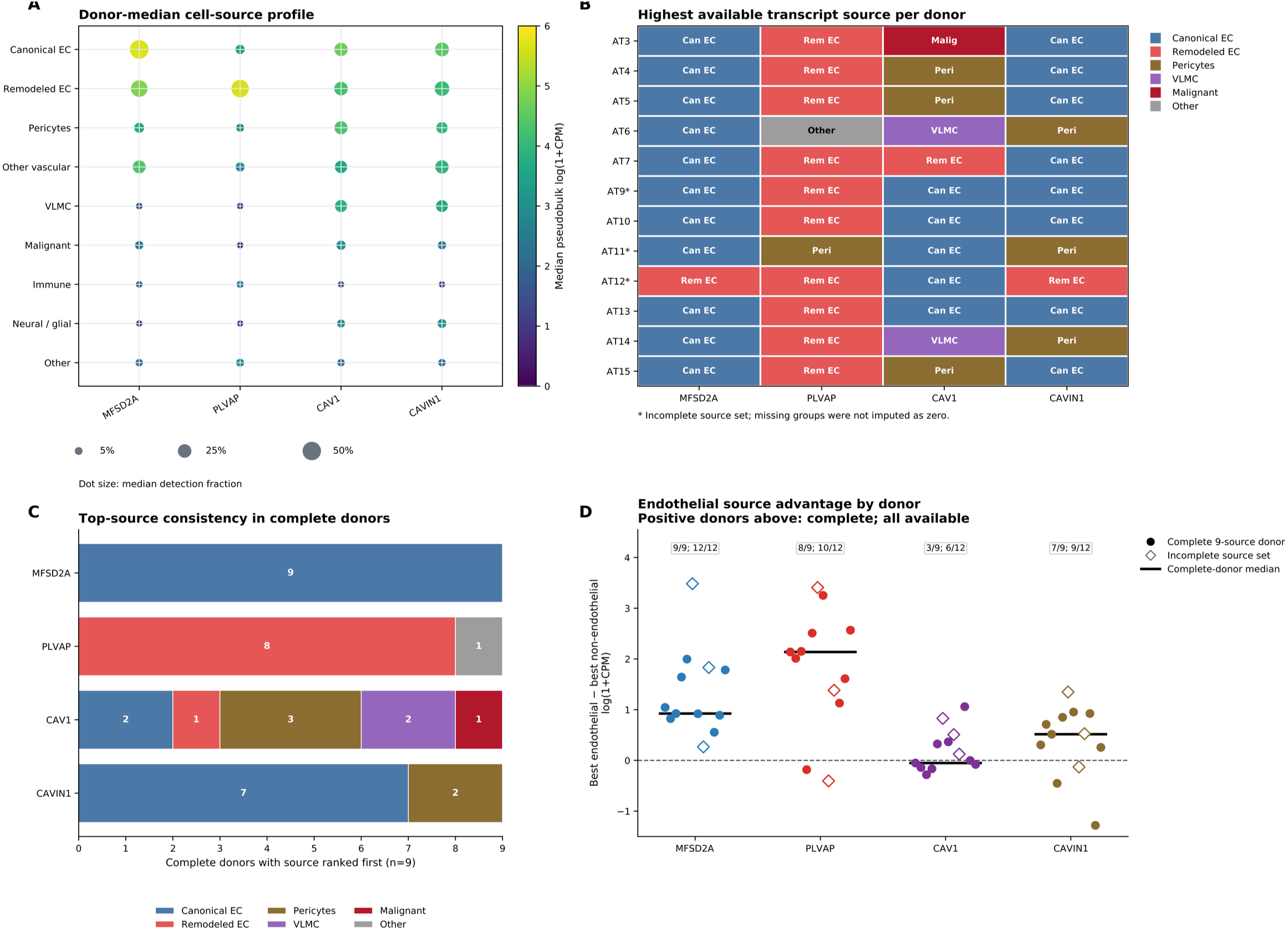
Donor-aware transcript-source localization across 9 predefined cellular source groups. (A) Donor-median profiles for *MFSD2A*, *PLVAP*, *CAV1*, and *CAVIN1*; dot size denotes median detection and color denotes median pseudobulk log(1+CPM). (B) Highest available source per donor; asterisks mark incomplete source sets, without zero imputation. (C) Top-source consistency among 9 complete donors. (D) Best endothelial-minus-non-endothelial advantage by donor; symbols distinguish complete and incomplete source sets, labels give positive-donor counts, and bars give complete-donor medians. Source localization does not establish protein localization, function, permeability, or causal remodeling. **Alt text:** Source-localization plots show *MFSD2A* in canonical endothelium, *PLVAP* in remodeled endothelium, and broader *CAV1*/*CAVIN1* vascular or mural sources.

### External datasets support spatial remodeling and malignant-state architecture

In GSE162631, all 4 paired patients showed lower BTB-integrity programs and higher vascular-remodeling, angiogenesis, and composite BTB-remodeling programs in tumor core versus peripheral tissue; barrier-dominant endothelial states were peripheral-enriched and remodeling/angiogenic endothelial states core-enriched in all 4 patients. In corrected Ivy ASTR122, vascular remodeling and the composite BTB-remodeling axis differed across LE, IT, and CT after Holm correction (*P* = 0.033327 and *P* = 0.018524, respectively), with the expected direction in 7/8 complete tumors for both regional contrasts. Xenium S2/S6 provided cross-technology localization support: among 106,778 source-annotated cells, including 18,337 strict endothelial cells, *PLVAP* and *CAV1* were endothelial-enriched; *THSD1*, *FLT4*, and *MFSD2A* were not measured (Fig. 5; Supplementary Fig. S5; Supplementary Table S5).

**Figure 5.**
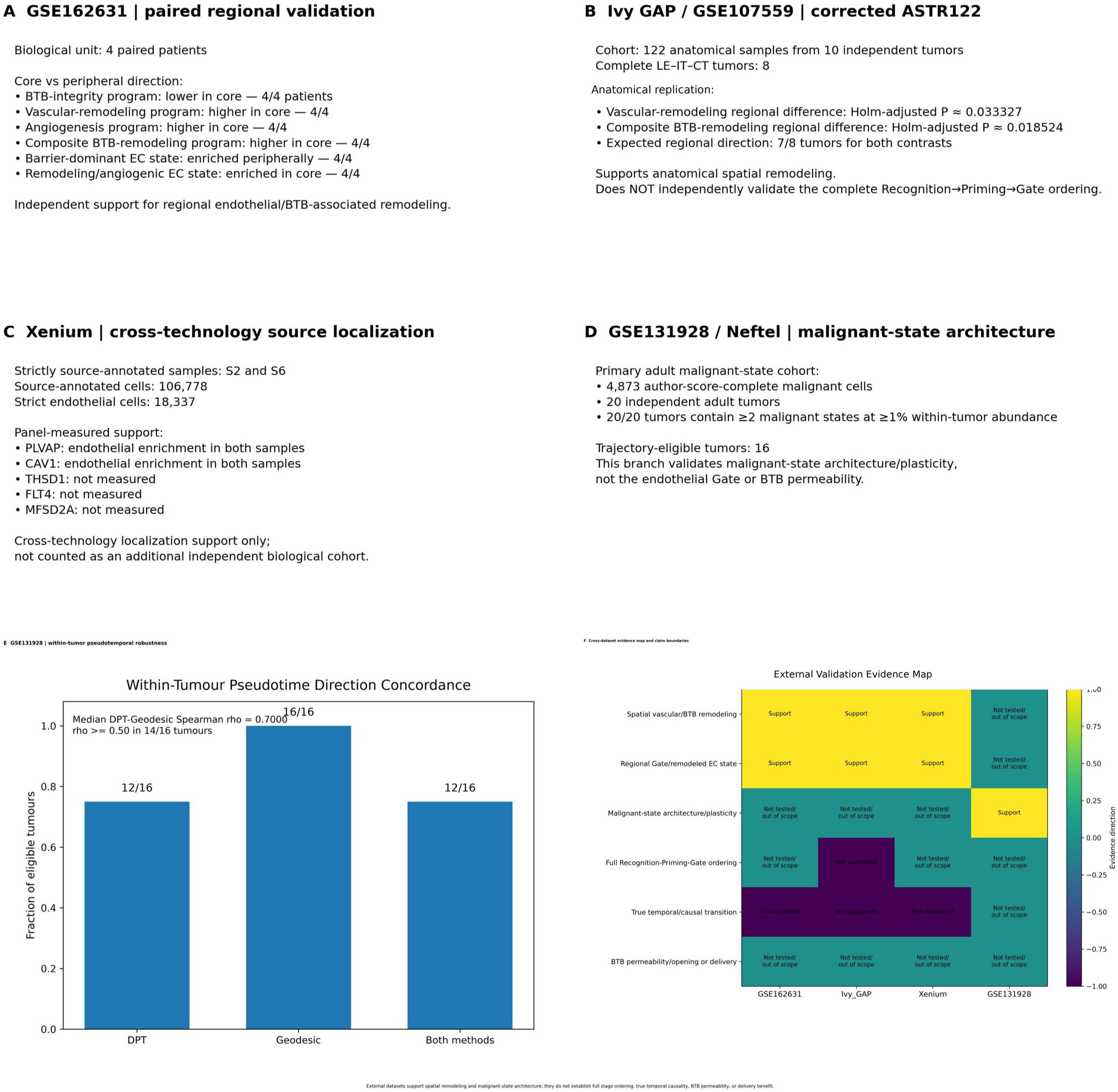
Complementary external validation of spatial vascular/BTB remodeling and malignant-state architecture. (A) GSE162631 patient-paired validation across 4 patients. (B) Corrected Ivy GAP ASTR122 anatomical replication across 10 tumors, including 8 complete LE/IT/CT tumors; vascular remodeling and composite BTB remodeling differed across regions after Holm correction. (C) Xenium S2/S6 cross-technology localization in 106,778 source-annotated cells, including 18,337 strict endothelial cells; *PLVAP* and *CAV1* were endothelial-enriched, while *THSD1*, *FLT4*, and *MFSD2A* were not measured. (D) GSE131928 malignant-state architecture in 4,873 malignant cells from 20 adult tumors. (E) Within-tumor pseudotemporal robustness in 16 eligible tumors. (F) Evidence map summarizing distinct validation roles and boundaries. External data do not independently validate a complete Recognition-Priming-Gate sequence, true temporal causality, measured BTB permeability, or therapeutic-delivery benefit. **Alt text:** External validation panels summarize paired regional remodeling, Ivy anatomical replication, Xenium localization, and malignant-state trajectory robustness.

### An independent adult cohort supports malignant-state plasticity

The primary GSE131928 analysis included 4,873 author-score-complete malignant cells from 20 adult tumors; all 20 contained at least two malignant states at a within-tumor abundance threshold of 1%. Sixteen tumors met trajectory-reconstruction criteria. AC/MES-associated cells occupied later positions than OPC/NPC-associated cells in 12/16 tumors by diffusion pseudotime and 16/16 by graph-geodesic distance; both measures were directionally concordant in 12/16 tumors. The median within-tumor Spearman correlation between trajectory measures was 0.7000, with *ρ* ≥ 0.50 in 14/16 tumors (Fig. 5; Supplementary Fig. S6; Supplementary Table S6).

### Integrated interpretation

Together, the validation layers converged on two reproducible but distinct axes. Patient-paired and anatomically resolved cohorts supported spatial vascular/BTB remodeling, whereas the independent adult malignant-cell cohort supported broad state architecture and tumor-level pseudotemporal organization. The data do not independently validate a complete Recognition-Priming-Gate sequence, true temporal lineage, measured BTB permeability/opening, or improved therapeutic delivery (Supplementary Fig. S7; Supplementary Tables S6-S7).

## Discussion

The principal finding is a spatially partitioned vascular/BTB-associated architecture rather than a uniform or monotonic barrier program. Recognition was most reproducible at the LE-to-IT boundary, Priming increased toward IT and then decreased toward CT, and Gate remodeling was strongest from IT to CT. Because these are cross-sectional anatomical contrasts, Recognition-Priming-Gate is best understood as model decomposition: it organizes separable regional behaviors without establishing that a cell or vessel traverses the states in time or that one module causes the next.

The second major finding is a cell-resolved endothelial remodeling axis. Higher *PLVAP* and lower *MFSD2A* in remodeled endothelium were reproducible across donors and were reinforced by donor-aware source localization. This convergence supports a *PLVAP*-high/*MFSD2A*-low endothelial transcriptional state associated with Gate remodeling. It does not demonstrate barrier opening. *PLVAP* and *MFSD2A* are relevant to endothelial barrier biology, but permeability, tracer passage, junctional function, and drug penetration were not measured directly.^1,2^

Malignant-state plasticity forms a parallel axis rather than a proven upstream driver of vascular remodeling. GSE131928 reproduced broad within-tumor OPC/NPC/AC/MES state architecture and supported an exploratory continuum from developmental-like OPC/NPC-associated states toward AC/MES-associated states.^3^ However, pseudotime describes geometry within a transcriptional manifold, not a time-lapse lineage. The current data therefore support coexistence of malignant-state plasticity and spatial vascular/BTB remodeling without establishing directional tumor-to-vessel causality.

The external-validation design deliberately assigned different questions to different datasets. Patient-paired GSE162631 supported core-versus-peripheral endothelial remodeling,^5^ Ivy GAP/GSE107559 provided tumor-level anatomical replication,^6^ Xenium supplied cross-technology transcript localization,^7^ and GSE131928 tested malignant-state architecture.^3^ This role separation avoids pooling incompatible biological units and clarifies which components are independently supported. The strongest external evidence concerns spatial vascular/BTB remodeling and malignant-state organization; evidence is weaker for ordering, interaction, and functional consequences.

Recent BTB-modulation studies place this spatial framework in a translational context. Digiovanni et al showed that patient-derived GBM differentiation state can alter endothelial barrier competence through IL-6/STAT3-dependent regulation of junctional and transporter programs, supporting the proposition that tumor state can actively influence barrier function.^8^ Jimenez-Macias et al identified a BTB transcriptional program including *CDH5* and showed that pharmacologic modulation can disrupt endothelial barrier properties, increase intratumoral cisplatin accumulation, and potentiate chemotherapy in GBM models.^9^ Cai et al demonstrated spatially targeted, reversible optical blood-brain-tumor barrier modulation that enhanced paclitaxel delivery and therapeutic efficacy in infiltrative and angiogenic GBM models.^10^ Together, these studies show that tumor state can regulate the barrier, BTB molecular programs can be perturbed, and transient barrier modulation can improve delivery.

Our study does not experimentally unite those three propositions. Its contribution is a donor-aware spatial map of where tumor-associated Recognition/Priming signals and endothelial Gate-associated remodeling coexist or segregate. If functionally validated, this spatial layer could help define anatomical niches and endothelial states for testing targeted, reversible BTB modulation. *THSD1*-*FLT4*, *PLVAP*/*MFSD2A*, IL-6/STAT3, *CDH5*-targeted modulation, and optical BTB modulation should not be treated as mechanistically equivalent.

Several design features strengthen the interpretation: biological inference was anchored to donors, patients, or tumors rather than nested observations; effect direction and biological-unit consistency were emphasized; external datasets were assigned non-overlapping roles; and transcript-source, spatial-state, malignant-state, and functional-barrier claims were kept separate. The main limitations are equally important. The design is cross-sectional, the number of independent spatial donors is modest, the full Recognition-Priming-Gate ordering lacks independent temporal validation, and no permeability, protein-level perturbation, or therapeutic-delivery experiment was performed. Xenium is a technical localization layer rather than an independent cohort, and its panel did not measure *THSD1*, *FLT4*, or *MFSD2A*.

In conclusion, the data support a spatially organized, coupled but non-causal dual-axis model of GBM biology: a vascular/BTB-associated remodeling axis centered on *PLVAP*-high/*MFSD2A*-low endothelial transcription and regionally structured Recognition/Priming/Gate contrasts, alongside a malignant-state plasticity axis. The framework generates specific predictions for functional and perturbational testing while preserving the evidentiary boundary that it does not yet establish temporal sequence, causal tumor-vessel signaling, functional BTB opening, or improved therapeutic delivery.

## Required Statements

### Ethics

This study involved secondary analysis of publicly available and/or de-identified datasets and did not involve new participant recruitment, intervention, or collection of identifiable human participant data. Ethics oversight and informed-consent procedures for the originating datasets are described in the corresponding source publications and data records. The originating GBM-Space study reports written informed consent and ethics approval by the Cambridge Local Research Ethics Committee (REC 18/EE/0172).

### Authorship

Conceptualization: Li Zheng, Lu Gan; Methodology: Li Zheng; Formal analysis: Li Zheng; Data curation: Li Zheng; Visualization: Li Zheng; Writing - original draft: Li Zheng; Writing - review & editing: Li Zheng, Lu Gan; Supervision: Lu Gan.

### Data Availability

All data analyzed in this study are third-party publicly available and/or de-identified datasets. Processed GBM-Space data are available from the GBM-Space portal (https://www.gbmspace.org/). The source-annotated Xenium S2 and S6 files used for cross-technology localization were obtained from the Zenodo dataset “10x Xenium data from: A spatially resolved human glioblastoma atlas reveals distinct cellular and molecular patterns of anatomical niches” (Zenodo record 17622242; DOI: 10.5281/zenodo.17622242), specifically the files “S2.tar pranali.gz” and “S6.tar Pranali.gz”. These files accompany the published source study by Sonpatki et al.^7^ GSE162631, GSE107559, and GSE131928 are available through NCBI Gene Expression Omnibus. Dataset-specific roles and provenance are summarized in Supplementary Table S5. No new participant-level dataset was generated. Analysis scripts underlying the reported results are available from the corresponding author upon reasonable request.

## References

1. Arvanitis CD, Ferraro GB, Jain RK. The blood-brain barrier and blood-tumour barrier in brain tumours and metastases. Nat Rev Cancer. 2020;20(1):26–41. doi:10.1038/s41568-019-0205-x.

2. Xie Y, Yang F, He L, et al. Single-cell dissection of the human blood-brain barrier and glioma blood-tumor barrier. Neuron. 2024;112(18):3089–3105.e7. doi:10.1016/j.neuron.2024.07.026.

3. Neftel C, Laffy J, Filbin MG, et al. An integrative model of cellular states, plasticity, and genetics for glioblastoma. Cell. 2019;178(4):835–849.e21. doi:10.1016/j.cell.2019.06.024.

4. de Jong G, Memi F, Gracia T, et al. A spatiotemporal cancer cell trajectory underlies glioblastoma heterogeneity. bioRxiv. 2025. doi:10.1101/2025.05.13.653495.

5. Xie Y, He L, Lugano R, et al. Key molecular alterations in endothelial cells in human glioblastoma uncovered through single-cell RNA sequencing. JCI Insight. 2021;6(15):e150861. doi:10.1172/jci.insight.150861.

6. Puchalski RB, Shah N, Miller J, et al. An anatomic transcriptional atlas of human glioblastoma. Science. 2018;360(6389):660–663. doi:10.1126/science.aaf2666.

7. Sonpatki P, Park HJ, Xing YL, et al. A spatially resolved human glioblastoma atlas reveals distinct cellular and molecular patterns of anatomical niches. Nat Commun. 2026;17:2951. doi:10.1038/s41467-026-69716-2.

8. Digiovanni S, Lorenzati M, Bianciotto OT, et al. Blood-brain barrier permeability increases with the differentiation of glioblastoma cells in vitro. Fluids Barriers CNS. 2024;21(1):89. doi:10.1186/s12987-024-00590-0.

9. Jimenez-Macias JL, Vaughn-Beaucaire P, Bharati A, et al. Modulation of blood-tumor barrier transcriptional programs improves intratumoral drug delivery and potentiates chemotherapy in GBM. Sci Adv. 2025;11(9):eadr1481. doi:10.1126/sciadv.adr1481.

10. Cai Q, Li X, Xiong H, et al. Optical blood-brain-tumor barrier modulation expands therapeutic options for glioblastoma treatment. Nat Commun. 2023;14(1):4934. doi:10.1038/s41467-023-40579-1.

