## Supplementary Material for "Spatial vascular/BTB remodeling and malignant-state plasticity in glioblastoma"

### Supplementary Methods

#### Patient- and donor-level robustness analysis

To make biological-unit replication explicit for the primary claims, prespecified summaries were generated for THSD1, FLT4, the THSD1–FLT4 Recognition score, Priming, Gate, PLVAP, and MFSD2A. For THSD1, FLT4, and Recognition, section-level adjusted regional effects were estimated by ordinary least squares with standardized vascular abundance and malignant fraction as covariates and then aggregated by the median within donor. Donor-level Priming/Gate and endothelial contrasts were reconstructed from the analysis-ready source tables and regression-checked against the reported primary results. Expected-direction counts were reported at the biological-unit level. Two-sided exact sign tests omitted exact zero effects and were interpreted descriptively. Median-effect uncertainty was summarized by 20,000 biological-unit bootstrap resamples using random seed 20260817. With four discovery donors, 4/4 direction consistency corresponds to a two-sided exact sign-test P=0.125 and was not treated as conventional statistical significance.

#### *THSD1–FLT4* spatial-proximity analysis

Spatial topology was evaluated in the same five-section Visium discovery cohort used for the primary LE–IT–CT analyses. Spatial coordinates were aligned to the primary spatial-location table, with all five sections passing exact spot-ID alignment. Within each section, *THSD1*-positive malignant-enriched spots were prespecified as source niches and *FLT4*-positive vascular-enriched spots as target niches. The primary malignant/vascular enrichment threshold was the within-section 50th percentile (q50), with q60 and q70 sensitivity analyses. A spatial-neighbor radius of 1.35 times the section-specific median first-nearest-neighbor distance was used. Same-spot co-occurrence, neighboring-target fraction, and nearest distinct target distance were evaluated. The primary q50 analysis used 5,000 permutations; q60/q70 sensitivity analyses used 2,000 permutations, with random seed 20260817. Source labels were resampled only among malignant-enriched spots within the same section while preserving source counts separately within LE, IT, and CT; *FLT4*-positive vascular target locations were held fixed. Section effects were aggregated by median within donor, so the two AT3 sections contributed one donor-level estimate. The analysis was interpreted only as spatial association and not as direct cell–cell contact, biochemical *THSD1–FLT4* signaling, temporal ordering, BTB permeability/opening, or therapeutic-delivery benefit.

#### Interpretation boundary

These supplementary robustness analyses do not revise the primary cohort definitions, reported primary effect estimates, or candidate definitions. The spatial-proximity analysis uses the same GBM-Space discovery cohort and is therefore a same-cohort spatial robustness analysis rather than independent external replication. All proximity findings are restricted to spatial association.

### Supplementary Figure Legends

Supplementary Figure S1. Cohort eligibility, quality control, and inferential hierarchy. (A) The discovery spatial cohort comprised 38 histopathology-annotated sections from six donors across leading-edge (LE), infiltrative-tumor (IT), cellular-tumor (CT), vascular (VASC), and necrotic (NEC) compartments. The LE–IT–CT ordering is anatomical and cross-sectional rather than temporal. (B) Eligibility was stable between MIN25 and MIN50; five complete LE–IT–CT sections from four donors were available at MIN50, and four complete sections from the same four donors remained at MIN100. (C) Donor/patient/tumor was treated as the biological inferential unit, with sections and spot-/cell-level observations nested within biological units. (D) Cohort counts used by primary spatial claims are summarized. Section-by-region and section-by-region-by-module records describe sampling structure and are not independent biological replicates.

**Supplementary Figure S2.** Threshold, pattern, and model robustness of spatial analyses. (A) *THSD1–FLT4* Recognition effects were compared between the MIN50 primary cohort and MIN100 sensitivity cohort; labels indicate the number of donors with positive effects. (B) Continuous Priming and Gate contrasts were similarly compared at MIN50 and MIN100. (C) Intended Priming LE–IT and Gate IT–CT directionality was retained across quantile thresholds from 0.50 to 0.70. (D) Fine vascular-composition CT–IT sensitivity models showed comparable directionality at MIN50 and MIN100. Fine-composition models are sensitivity analyses only; early IT–LE detailed models exhibited collinearity and are not treated as primary evidence.

Supplementary Figure S3. Patient/donor-level robustness of primary claims. (A) Donor-level spatial effects for THSD1, FLT4, the THSD1–FLT4 Recognition score, Priming, and Gate. (B) Donor-level remodeled-versus-canonical endothelial pseudobulk effects for PLVAP and MFSD2A. (C) Fraction of biological units in the prespecified expected direction. THSD1, Recognition, and Gate showed 4/4 donor directional consistency in the spatial discovery cohort; PLVAP and MFSD2A showed strong donor-level directionality in the 12-donor endothelial layer. Exact sign-test P values are descriptive; with n=4, 4/4 directional consistency corresponds to a two-sided exact sign-test P=0.125.

**Supplementary Figure S4.** *THSD1–FLT4* spatial-proximity robustness in the five-section Visium discovery cohort. *THSD1*-positive malignant-enriched spots were prespecified as source niches and *FLT4*-positive vascular-enriched spots as target niches. (A) Neighborhood enrichment relative to within-section, LE/IT/CT-stratified permutation expectation. (B) Nearest-target closeness, defined as permutation-median minus observed nearest distinct target distance in normalized nearest-neighbor units. (C) Same-spot co-occurrence relative to permutation expectation. (D) Donor-level directional consistency across q50, q60, and q70 compartment-enrichment thresholds. AT3 sections were aggregated before donor-level interpretation. The analysis supports spatial association only; Visium spots are multicellular and do not establish direct tumor–endothelial contact, biochemical *THSD1–FLT4* signaling, or BTB permeability.

**Supplementary Figure S5.** Complementary external validation and evidence boundaries. (A) Roles of GSE162631, corrected Ivy GAP/GSE107559 ASTR122, Xenium, and GSE131928/Neftel in the external-validation framework. (B) GSE162631 showed concordant core-versus-peripheral endothelial/BTB-associated remodeling in 4/4 paired patients; corrected Ivy ASTR122 supported anatomical vascular and composite BTB-remodeling differences; Xenium provided cross-technology source-localization support but did not measure *THSD1*, *FLT4*, or *MFSD2A*. (C) Cross-dataset evidence map. These datasets support spatial remodeling and malignant-state architecture but do not independently validate the complete Recognition → Priming → Gate ordering, true temporal/causal transition, measured BTB permeability/opening, or delivery benefit.

**Supplementary Figure S6.** Independent malignant-state architecture and pseudotemporal robustness in GSE131928/Neftel. (A) The primary adult malignant-state cohort comprised 4,873 author-score-complete malignant cells from 20 independent adult tumors; all 20 tumors contained at least two malignant states at ≥1% within-tumor abundance. (B) Sixteen tumors passed within-tumor pseudotemporal reconstruction quality control; diffusion pseudotime showed the expected direction in 12/16 tumors, graph-geodesic ordering in 16/16, and both methods were positive in 12/16. (C) Direction concordance and agreement between trajectory measures. This branch supports malignant-state architecture/plasticity and an exploratory within-tumor continuum; it does not establish true temporal lineage or validate the endothelial Gate.

Supplementary Figure S7. Claim boundaries, dataset roles, and non-equivalences. (A) Claim-level evidence status and the corresponding interpretation boundary. (B) Dataset-role rules distinguish donor-aware analyses within the same GBM-Space atlas from independent external biological replication, and distinguish Xenium cross-technology source localization from an independent cohort. (C) Interpretation boundaries identify inferences not established by the present study, including validated temporal Recognition→Priming→Gate sequencing, biochemical THSD1→FLT4 activation, equivalence of transcript states with BTB permeability/opening, and improved therapeutic delivery. These boundaries are retained regardless of directional consistency in computational analyses.

### Supplementary Tables

#### Supplementary Table S1. Discovery cohort, eligibility, quality control, and biological inferential-unit summary.

| Item | Value | Interpretation |
|---|---|---|
| Discovery spatial cohort | 38 sections / 6 donors | Histopathology-annotated GBM-Space discovery cohort. |
| Anatomical compartments | LE; IT; CT; VASC; NEC | Cross-sectional anatomical compartments. |
| IT–CT MIN50 eligibility | 17 sections / 4 donors | Used for IT–CT analyses. |
| Complete LE–IT–CT MIN50 | 5 sections / 4 donors | Primary complete three-compartment spatial cohort. |
| Complete LE–IT–CT MIN100 | 4 sections / 4 donors | Sensitivity cohort; same four donors. |
| MIN25 vs MIN50 | Qualification counts identical | No expansion of biological donor count. |
| Biological inferential unit | donor / patient / tumor | Sections, spots, nuclei and cells are nested observations. |
| Section-by-region records | 190 | Sampling records; not independent biological n. |
| Section-by-region-by-module records | 135 | Sampling records; not independent biological n. |

**Supplementary Table S2. MIN50-versus-MIN100 and model-robustness summary for Recognition, Priming, Gate, and vascular-composition sensitivity analyses.**

| Layer | Metric | MIN50 median effect | MIN50 direction | MIN100 median effect | MIN100 direction | Interpretation |
|---|---|---|---|---|---|---|
| Recognition | Recognition LE→IT | 0.0287 | 4/4 positive | 0.0287 | 4/4 positive | Adjusted donor-level spatial effect |
| Recognition | Recognition IT→CT | -0.0162 | 2/4 positive | -0.0162 | 2/4 positive | Adjusted donor-level spatial effect |
| Priming | Priming LE→IT | 0.1481 | 3/4 positive | 0.2089 | 3/4 positive | Continuous-score donor contrast |
| Priming | Priming IT→CT | -0.1641 | 0/4 positive | -0.2103 | 0/4 positive | Continuous-score donor contrast |
| Gate | Gate LE→IT | -0.0641 | 1/4 positive | -0.1663 | 1/4 positive | Continuous-score donor contrast |
| Gate | Gate IT→CT | 0.2130 | 4/4 positive | 0.2333 | 4/4 positive | Continuous-score donor contrast |
| Vascular composition sensitivity | *CAV1* CT→IT detailed β | 0.2753 | 4/4 positive | 0.2753 | 4/4 positive | Sensitivity model; not stronger than primary donor-level evidence |
| Vascular composition sensitivity | *CAVIN1* CT→IT detailed β | 0.2044 | 4/4 positive | 0.2044 | 3/4 positive | Sensitivity model; not stronger than primary donor-level evidence |
| Vascular composition sensitivity | *MFSD2A* CT→IT detailed β | 0.2150 | 4/4 positive | 0.1875 | 4/4 positive | Sensitivity model; not stronger than primary donor-level evidence |
| Vascular composition sensitivity | *PLVAP* CT→IT detailed β | 0.1053 | 3/4 positive | 0.0731 | 3/4 positive | Sensitivity model; not stronger than primary donor-level evidence |

### Supplementary Table S3. Biological-unit robustness summary for primary spatial and endothelial claims.

| Signal | Contrast | Biological units | Median effect | Bootstrap 95% interval | Expected direction | Units in expected direction | Exact two-sided sign-test P | Interpretation |
|---|---|---|---|---|---|---|---|---|
| *FLT4* | LE→IT | 4 | 0.0820 | -0.1485 to 0.6028 | positive | 3/4 | 0.6250 | Biological-unit inference; sign test descriptive. |
| Gate | IT→CT | 4 | 0.2130 | 0.1449 to 0.5392 | positive | 4/4 | 0.1250 | Biological-unit inference; sign test descriptive. |
| *MFSD2A* | remodeled−canonical | 12 | -0.7806 | -1.1427 to -0.3782 | negative | 10/12 | 0.0386 | Biological-unit inference; sign test descriptive. |
| *PLVAP* | remodeled−canonical | 12 | 1.8995 | 1.2311 to 2.1644 | positive | 11/12 | 0.0063 | Biological-unit inference; sign test descriptive. |
| Priming | LE→IT | 4 | 0.1481 | -0.0107 to 0.3068 | positive | 3/4 | 0.6250 | Biological-unit inference; sign test descriptive. |
| *THSD1–FLT4* Recognition | LE→IT | 4 | 0.0287 | 0.0082 to 0.3541 | positive | 4/4 | 0.1250 | Biological-unit inference; sign test descriptive. |
| *THSD1* | LE→IT | 4 | 0.2175 | 0.1678 to 0.3431 | positive | 4/4 | 0.1250 | Biological-unit inference; sign test descriptive. |

### Supplementary Table S4. *THSD1–FLT4* spatial-proximity results and threshold sensitivity.

| Analysis | Metric | Threshold | Donors | Positive donors | Median donor effect | Interpretation |
|---|---|---|---|---|---|---|
| Primary donor direction | Same-spot co-occurrence | q50 (primary) | 4 | 3/4 | 0.0104 | Positive values indicate stronger *THSD1*-malignant/*FLT4*-vascular spatial association than the region-stratified null. |
| Primary donor direction | Neighborhood enrichment | q50 (primary) | 4 | 4/4 | 0.0079 | Positive values indicate stronger *THSD1*-malignant/*FLT4*-vascular spatial association than the region-stratified null. |
| Primary donor direction | Nearest-target closeness | q50 (primary) | 4 | 3/4 | 0.7326 | Positive values indicate stronger *THSD1*-malignant/*FLT4*-vascular spatial association than the region-stratified null. |
| Threshold sensitivity | Same-spot co-occurrence | q50 | 4 | 3/4 | 0.0104 | Direction sensitivity across prespecified enrichment thresholds. |
| Threshold sensitivity | Neighborhood enrichment | q50 | 4 | 4/4 | 0.0079 | Direction sensitivity across prespecified enrichment thresholds. |
| Threshold sensitivity | Nearest-target closeness | q50 | 4 | 3/4 | 0.7326 | Direction sensitivity across prespecified enrichment thresholds. |
| Threshold sensitivity | Same-spot co-occurrence | q60 | 4 | 3/4 | 0.0090 | Direction sensitivity across prespecified enrichment thresholds. |
| Threshold sensitivity | Neighborhood enrichment | q60 | 4 | 4/4 | 0.0072 | Direction sensitivity across prespecified enrichment thresholds. |
| Threshold sensitivity | Nearest-target closeness | q60 | 4 | 3/4 | 0.4146 | Direction sensitivity across prespecified enrichment thresholds. |
| Threshold sensitivity | Same-spot co-occurrence | q70 | 4 | 3/4 | 0.0075 | Direction sensitivity across prespecified enrichment thresholds. |
| Threshold sensitivity | Neighborhood enrichment | q70 | 4 | 4/4 | 0.0064 | Direction sensitivity across prespecified enrichment thresholds. |
| Threshold sensitivity | Nearest-target closeness | q70 | 4 | 3/4 | 0.5302 | Direction sensitivity across prespecified enrichment thresholds. |

### Supplementary Table S5. External-validation datasets, biological units, roles, and interpretation boundaries.

| Dataset | Biological unit | Role | Key result | Supports | Does not establish |
|---|---|---|---|---|---|
| GSE162631 | 4 paired patients | Independent patient-paired regional endothelial remodeling validation | Core<Peripheral BTB integrity 4/4; Core>Peripheral vascular remodeling 4/4; Core>Peripheral angiogenesis 4/4; Core>Peripheral composite BTB remodeling 4/4; barrier-dominant EC Peripheral>Core 4/4; remodeling/angiogenic EC Core>Peripheral 4/4 | Spatial endothelial remodeling axis; regionally structured BTB state | True temporal lineage; measured permeability; causality |
| Ivy GAP / GSE107559 ASTR122 | 8 complete LE+IT+CT tumors | Independent anatomical replication of vascular/BTB remodeling programs | Vascular remodeling Friedman Holm P=0.033327; BTB remodeling axis Friedman Holm P=0.018524 | Anatomical spatial vascular remodeling; BTB remodeling axis | Full Recognition–Priming–Gate stage ordering; individual-gene stage localization; monotonic BTB-integrity loss; permeability; causality |
| Xenium S2/S6 | 2 source-annotated samples: S2 and S6 | Cross-technology transcript-source localization. Source: Sonpatki et al., Nat Commun. 2026;17:2951 (doi:10.1038/s41467-026-69716-2); Zenodo 17622242 (doi:10.5281/zenodo.17622242), files “S2.tar pranali.gz” and “S6.tar Pranali.gz”. | 106,778 exact source-annotated cells; 18,337 strict endothelial cells; *PLVAP* and *CAV1* higher in strict endothelial vs nonvascular in S2/S6 across detection, mean count and panel-CP10K | *PLVAP* endothelial source localization; descriptive *CAV1* endothelial enrichment | Independent cohort replication; negative conclusions for *THSD1*, *FLT4*, or *MFSD2A*; *CAV1* endothelial specificity; permeability; causality |
| GSE131928 / Neftel Smart-seq2 | 20 adult tumors; 16 eligible within-tumour trajectories | Independent malignant-state architecture and exploratory within-tumor continuum | 4,873 author-score-complete adult malignant cells / 20 tumors; within-tumour trajectories PASS 16/16; DPT direction 12/16; geodesic direction 16/16; both methods 12/16; median DPT-geodesic rho 0.7000 | Adult malignant-state architecture; exploratory within-tumor pseudotemporal continuum | Direct precedence of Priming relative to vascular remodeling; true temporal lineage; endothelial Gate state; BTB permeability; causality |

### Supplementary Table S6. Cross-dataset support matrix for manuscript-level claims.

| **Claim** | **GSE162631** | **Ivy GAP/GSE107559** | **Xenium** | **GSE131928** | **Permitted manuscript interpretation** |
|---|---|---|---|---|---|
| Spatial vascular/BTB remodeling axis | Strong support | Strong support | Source-localization support | Out of scope | Independent patient-paired endothelial and anatomically resolved cohorts support spatial vascular/BTB remodeling associated with GBM anatomical compartment. |
| Regionally structured Gate/remodeled endothelial state | Strong program-level support | Spatial-remodeling support; not stage ordering | *PLVAP*/*CAV1* localization support | Out of scope | Core-associated regions show concordant endothelial/vascular remodeling and regionally structured BTB-state differences. |
| Malignant-state architecture/plasticity continuum | Out of scope | Out of scope | Out of scope | Strong exploratory support | An independent adult GBM cohort supports broad malignant-state architecture and a strong exploratory within-tumour pseudotemporal continuum from OPC/NPC-associated toward AC/MES-associated states. |
| Full Recognition -> Priming -> Gate stage ordering | Full sequence not tested | Not independently replicated | Not tested | Not tested | The full staged Recognition-Priming-Gate sequence remains a discovery-model interpretation and is not independently validated. |
| True temporal/causal transition | Not supported | Not supported | Not supported | Exploratory pseudotime only | Cross-sectional and pseudotemporal patterns do not establish true temporal lineage or causal conversion. |
| Measured BTB permeability/opening or improved delivery | Not tested | Not tested | Not tested | Not tested | No external-validation layer directly measures BTB permeability, opening, or therapeutic delivery. |

### Supplementary Table S7. Claim-level evidence status and interpretation boundaries.

| Claim | Evidence status | Evidence | Boundary |
|---|---|---|---|
| Spatial vascular/BTB remodeling axis | Strong Independent External Support | Discovery spatial cohort + GSE162631 + Ivy ASTR122 | Transcript/program-level spatial remodeling; not measured permeability. |
| *PLVAP*-high / *MFSD2A*-low remodeled endothelial transcriptional axis | Donor-Aware Transcript-Level Support | 12-donor GBM-Space single-nucleus endothelial layer | Same GBM-Space atlas, separately analysed; not protein function or BTB opening. |
| Malignant-state architecture/plasticity continuum | Strong External Malignant-State Support | GSE131928/Neftel adult malignant cohort | Pseudotime remains exploratory; not true lineage. |
| Recognition / Priming / Gate spatial organization | Discovery Model With Partial External Support | LE–IT–CT discovery + external remodeling/state evidence | Do not state validated temporal sequence or causality. |
| Full Recognition → Priming → Gate stage ordering | Not Independently Validated | Not replicated as full sequence | Discovery-model interpretation only. |
| Measured BTB permeability/opening or improved delivery | Not tested / not established | No current dataset directly measures this endpoint | No current dataset directly measures this endpoint. |
| *THSD1*-positive malignant-enriched spots are spatially associated with *FLT4*-positive vascular-enriched spots | Allowed As Spatial Association | *THSD1–FLT4* spatial-proximity analysis | Allowed only as a spatial-association statement when donor-level neighborhood/distance results support it. |
| *THSD1*-positive tumor cells directly contact *FLT4*-positive endothelial cells | Not established by present study | *THSD1–FLT4* spatial-proximity analysis | Visium spots are multicellular; direct cell-cell contact is not established. |
| *THSD1* activates *FLT4* | Not established by present study | *THSD1–FLT4* spatial-proximity analysis | No biochemical interaction or causal signaling experiment is performed. |
| *THSD1–FLT4* is a validated biochemical Recognition gate | Not established by present study | *THSD1–FLT4* spatial-proximity analysis | Recognition remains a cross-sectional spatial discovery framework. |
| The spatial association demonstrates BTB opening/permeability | Not established by present study | *THSD1–FLT4* spatial-proximity analysis | No permeability, transvascular transport or therapeutic-delivery measurement. |
| LE → IT → CT is a temporal trajectory | Not established by present study | *THSD1–FLT4* spatial-proximity analysis | LE/IT/CT remain cross-sectional anatomical compartments. |
