## Supplementary figures and images for "Spatial vascular/BTB remodeling and malignant-state plasticity in glioblastoma"

### Supplementary Figure S1

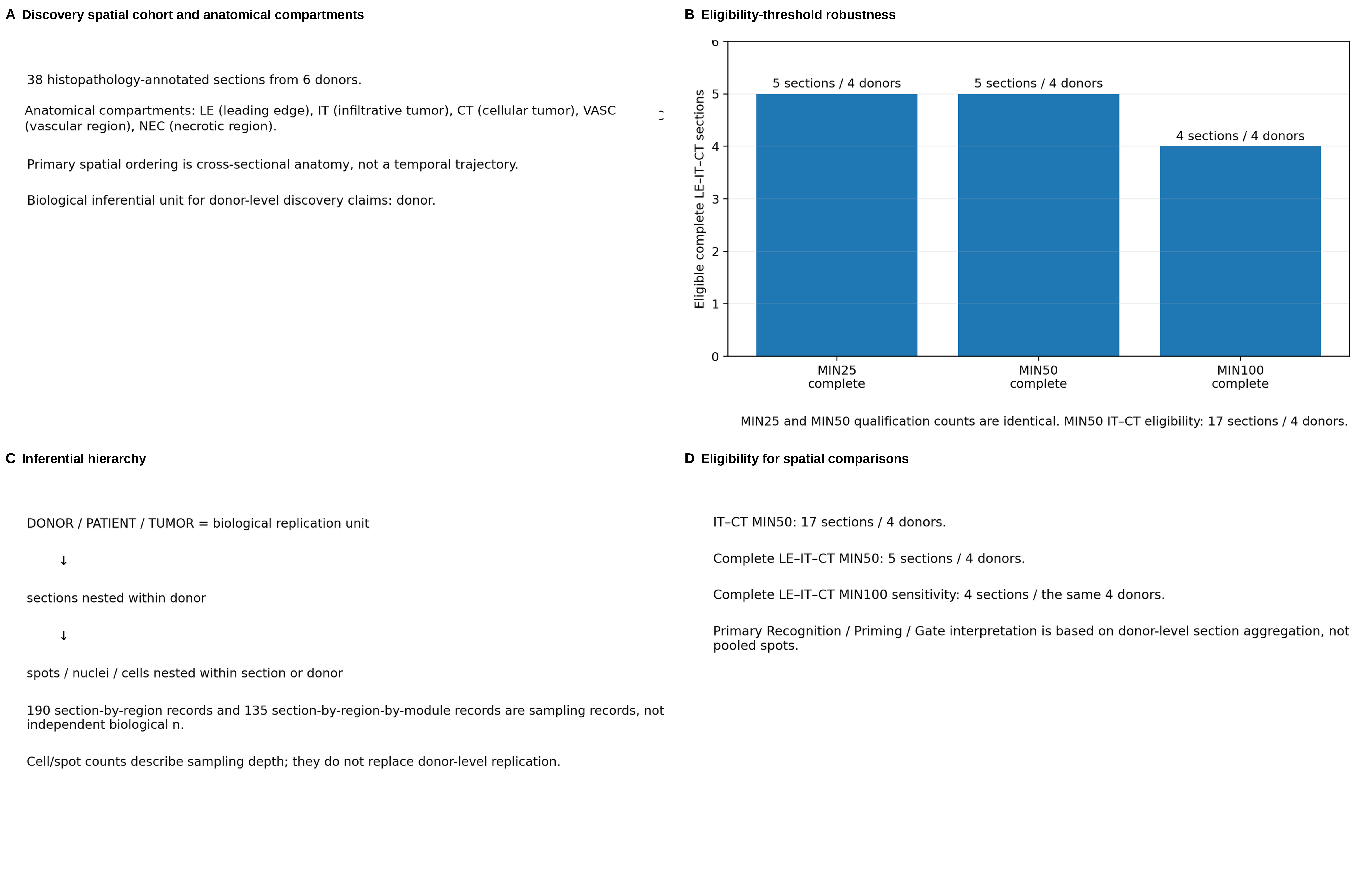

### Supplementary Figure S2

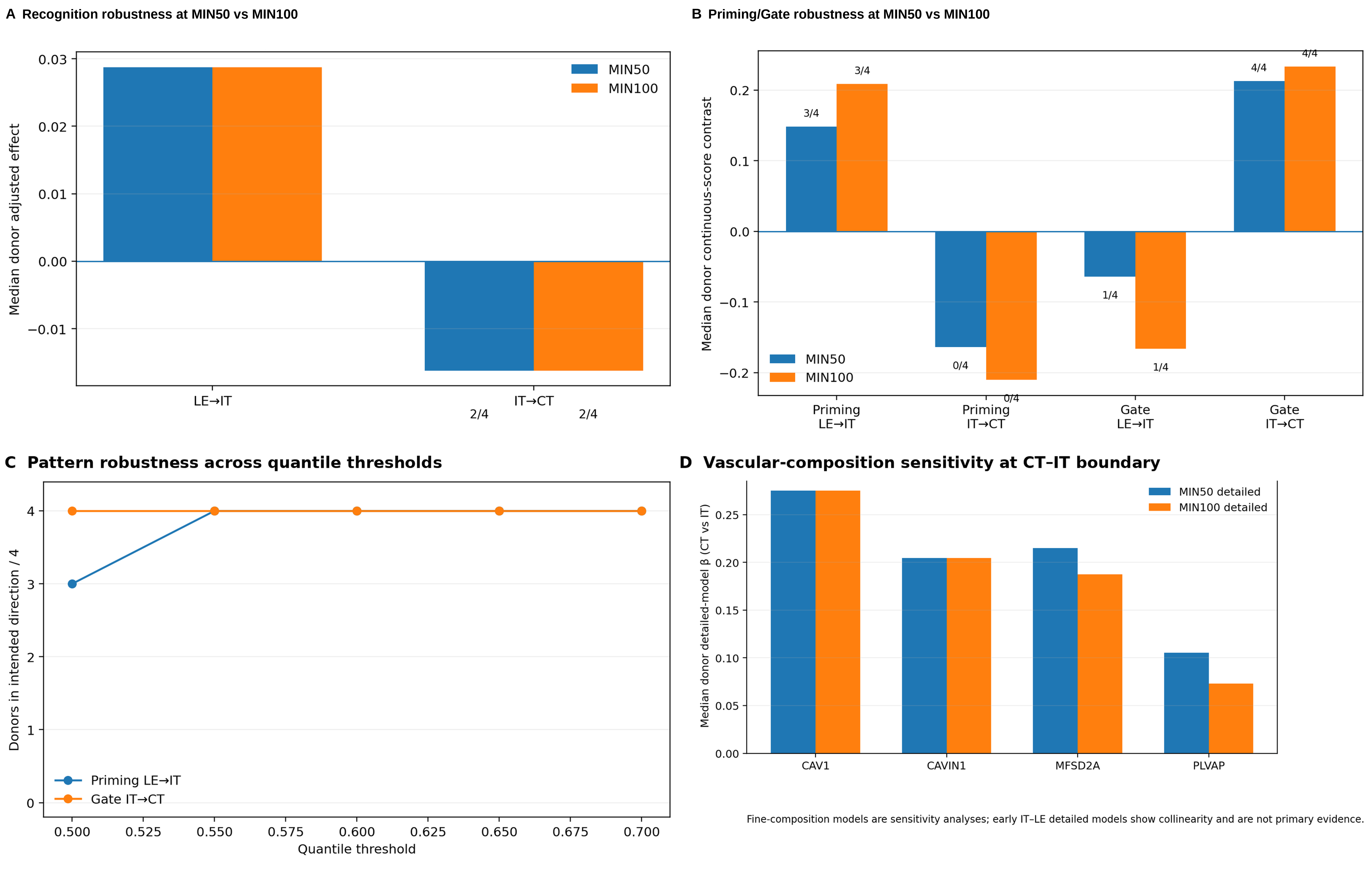

### Supplementary Figure S3

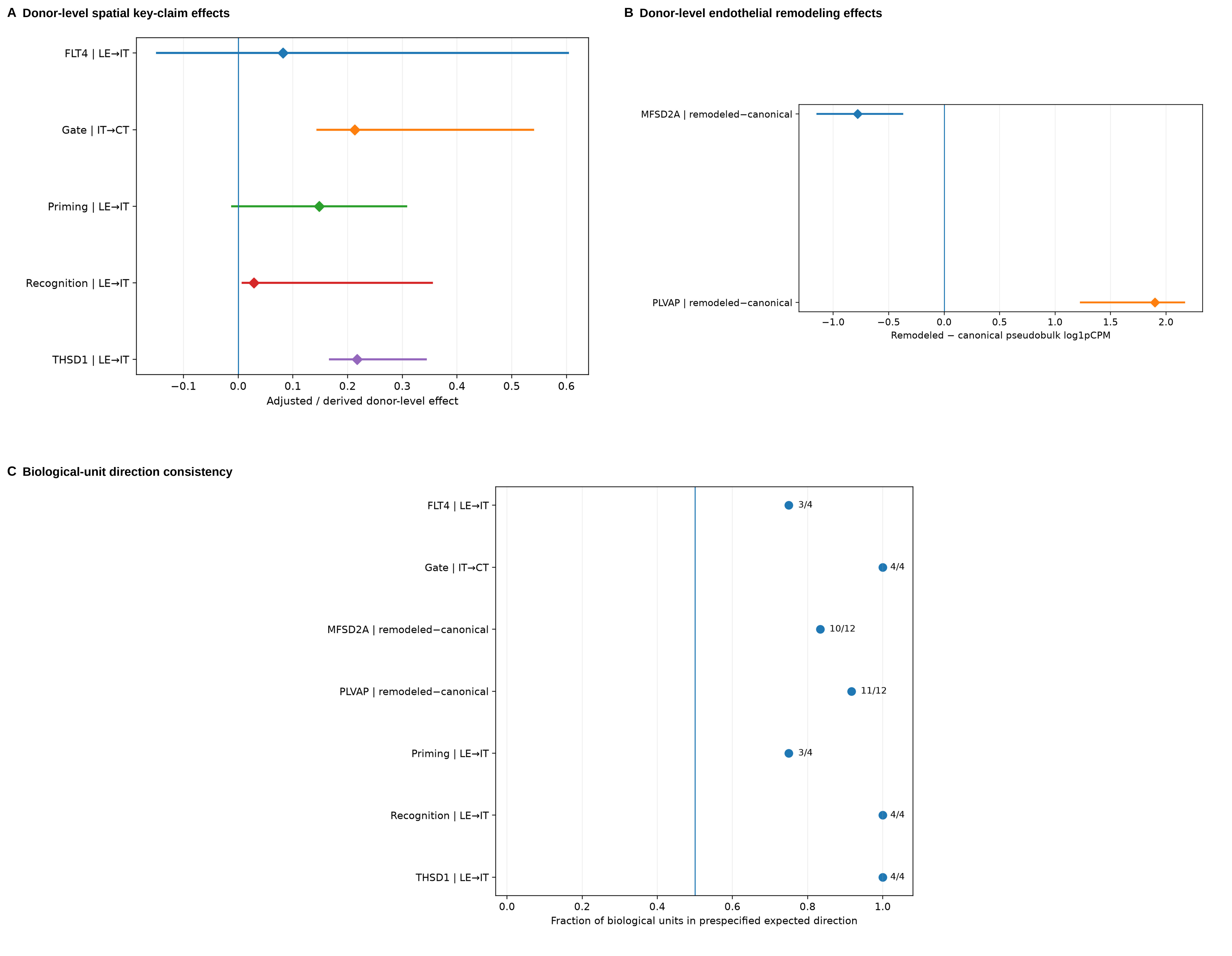

### Supplementary Figure S4

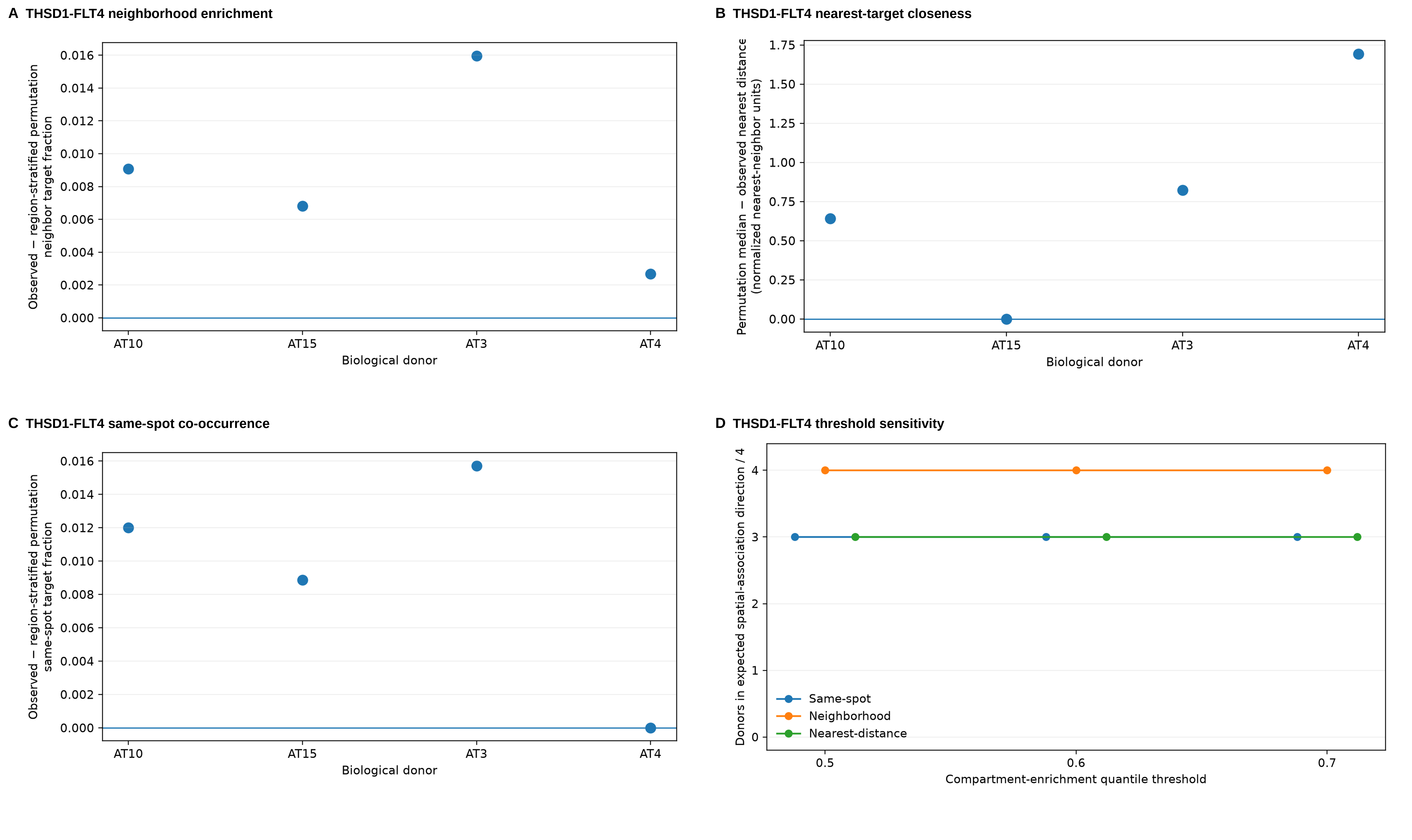

### Supplementary Figure S5

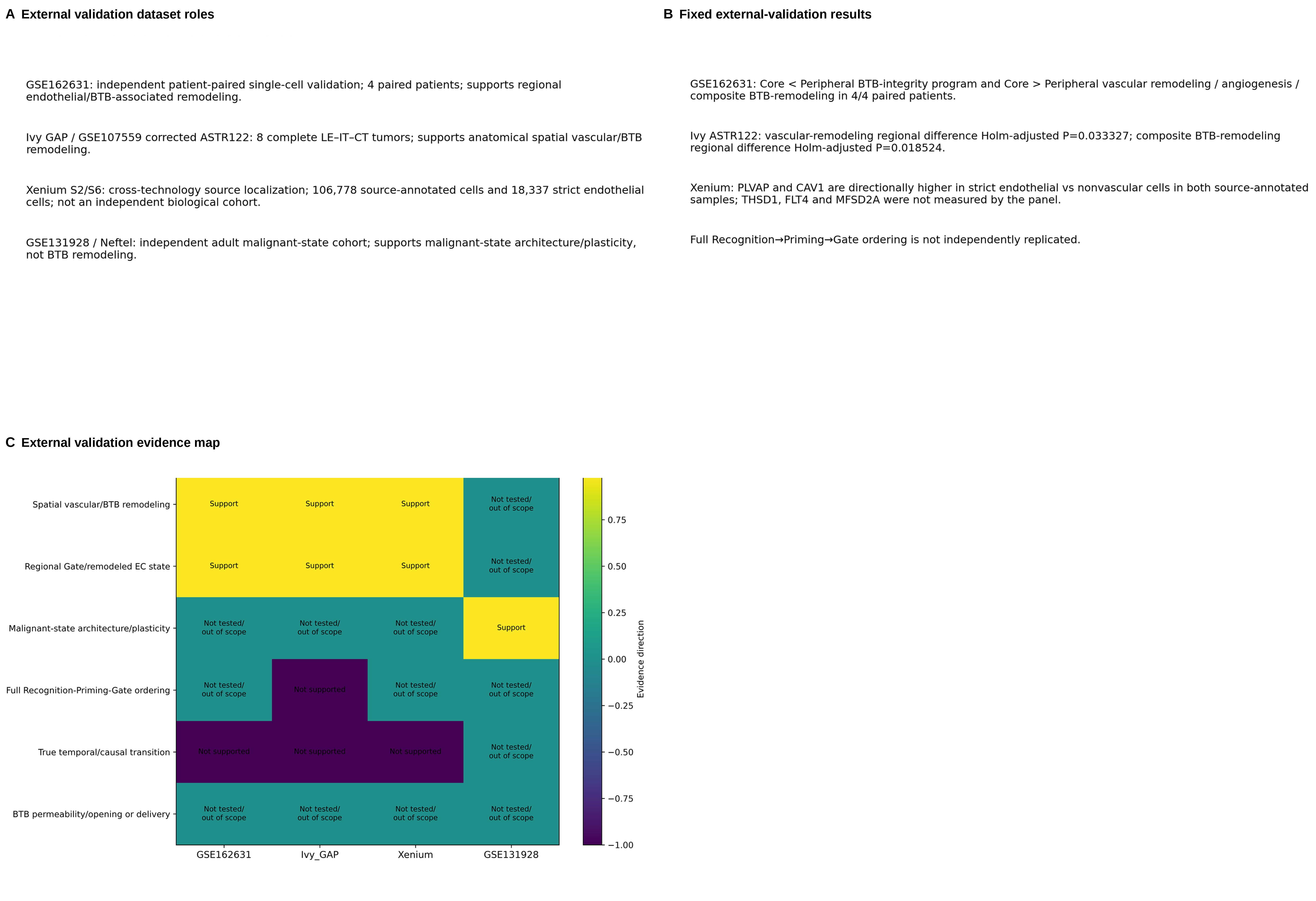

### Supplementary Figure S6

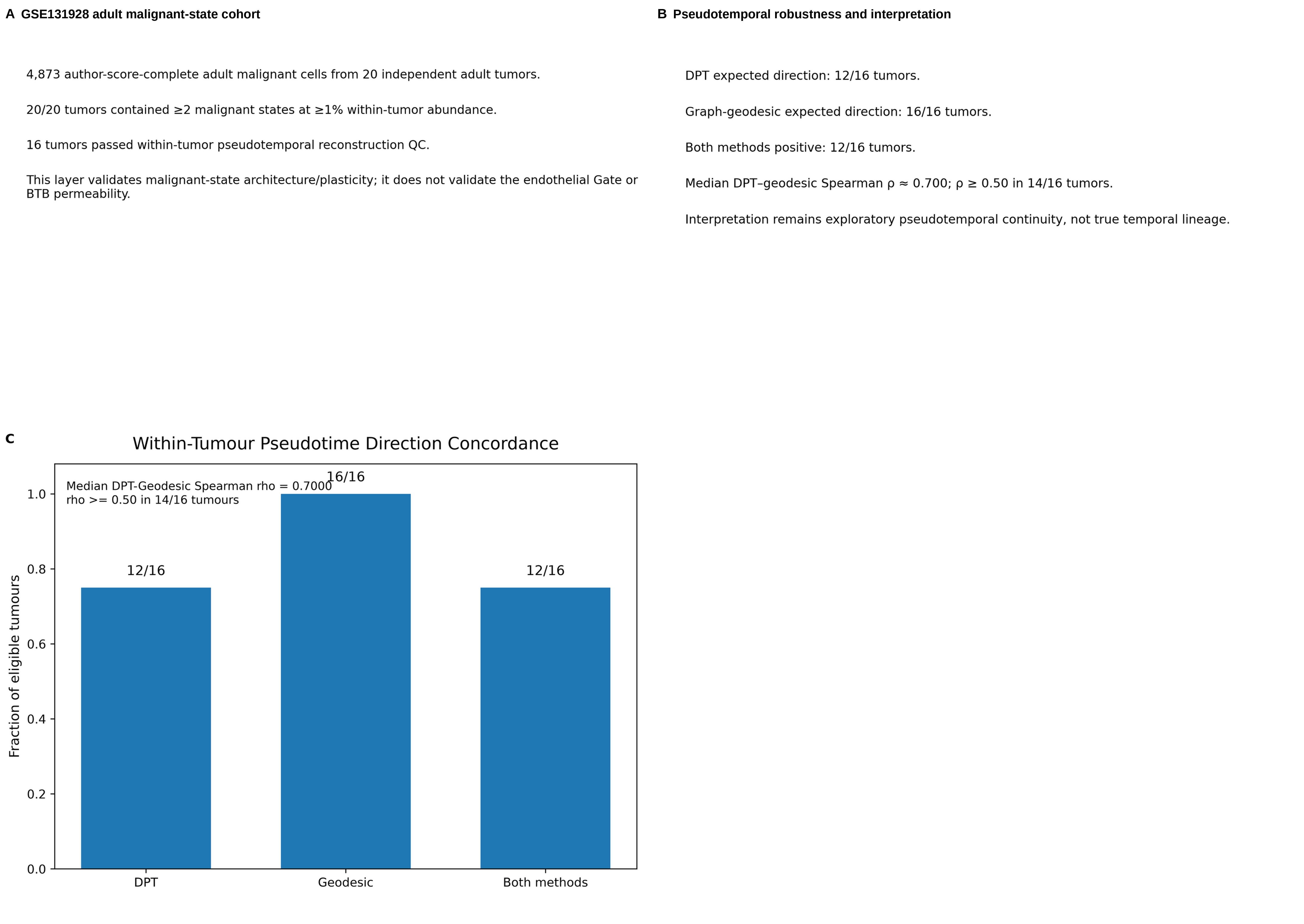

### Supplementary Figure S7

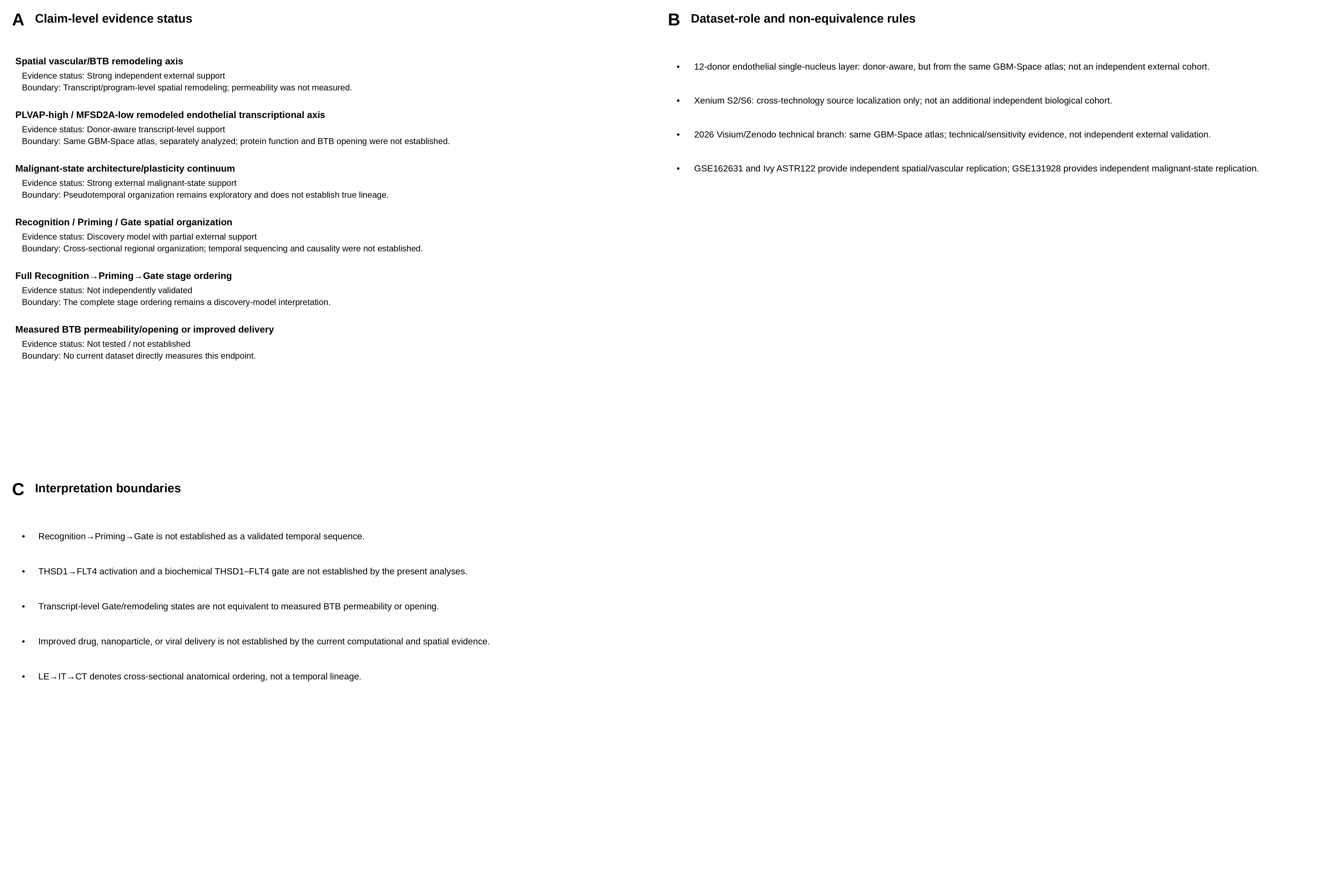
